# From Retrospective to Prospective: Integrated Value Representation in Frontal Cortex for Predictive Choice Behavior

**DOI:** 10.1101/2021.12.27.474215

**Authors:** Kosuke Hamaguchi, Hiromi Takahashi-Aoki, Dai Watanabe

**Affiliations:** Department of Biological Sciences, Kyoto University Graduate School of Medicine, Kyoto, Japan; Graduate School of Brain Science, Doshisha University, Kyoto, Japan

## Abstract

Animals must flexibly estimate the value of their actions to successfully adapt in a changing environment. The brain is thought to estimate action-value from two different sources, namely the action-outcome history (retrospective value) and the knowledge of the environment (prospective value). How these two different estimates of action-value are reconciled to make a choice is not well understood. Here we show that as a mouse learns the state-transition structure of a decision-making task, retrospective and prospective values become jointly encoded in the preparatory activity of neurons in the frontal cortex. Suppressing this preparatory activity in expert mice returned their behavior to a naïve state. These results reveal the neural circuit that integrates knowledge about the past and future to support predictive decision-making.

**One-Sentence Summary:** Preparatory activity in the mouse frontal cortex encodes prospective value to support predictive choice behavior.

## Main Text

An adaptive animal in a changing environment must flexibly change their choices, which depends on the ability to assess the value of each action (action-value). In a novel environment, a naïve animal would assess the action-value from the history of actions and outcomes (retrospective value). Computationally, the retrospective value can be calculated as the discounted sum of past rewards (*1, 2*) and is detected in various brain regions (*3-5*). However, the choice strategies based on the retrospective value can be slow, especially in a changing environment, as it requires trial-and-errors to update the value. By contrast, after learning the model of the environment, the action-value computed from the knowledge of the environment (prospective value) allows the animal to predict the state transition and promotes flexible and predictive choices (*6, 7*). Numerous brain regions important for the choice behavior that depends on the knowledge about the state transition within a trial (*8-10*), and over trials (e.g., change of reward condition) (*11-13*) were identified. Accumulating evidence suggests that retrospective and prospective values are represented in different cortico-striatal circuits; the posterior regions of the cortex and striatum are implicated in retrospective valuation and the regions of prefrontal cortex and anterior striatum in prospective valuation (*8, 14-16*). The fundamental yet unanswered question is how the brain selects the optimal action when different brain regions propose different values for the same action.

A subregion of the mouse frontal cortex, anterior lateral motor (ALM) cortex (*17*), is involved in the control and planning of orofacial movements (*18-20*). Higher-order motor cortices including ALM are also known to encode value-related information (*5, 21-23*). However, it is not known whether ALM encodes either retrospective or prospective values, or integrates both to make a choice.

To study the neuronal process to integrate retrospective and prospective values in mice, we designed a novel two-alternative forced-choice task. In this task, the reward condition (state) deterministically changes after every tenth reward delivery (Fig. 1A and B, we term “Reward10” task). This deterministic task structure allows the analytical calculation of the prospective value without relying on algorithms. Here, the prospective value is formulated as the discounted sum of future rewards (*1, 7*) (Methods). In this task, each trial begins with a preparatory period signaled by an auditory Ready signal (noise) during which the head-fixed mice (n = 43) were required to withhold licking. Upon hearing the Go tone, the mice were allowed to make one response (lick toward the left or right water port). In Left (Right) state, only the response toward left (right) is rewarded. The next trial resumes after an inter-trial interval (ITI, ∼10 s). After mice harvested all the rewards, the rewarding port was reversed without notice (Fig. 1B, state transition).

**Figure 1.**
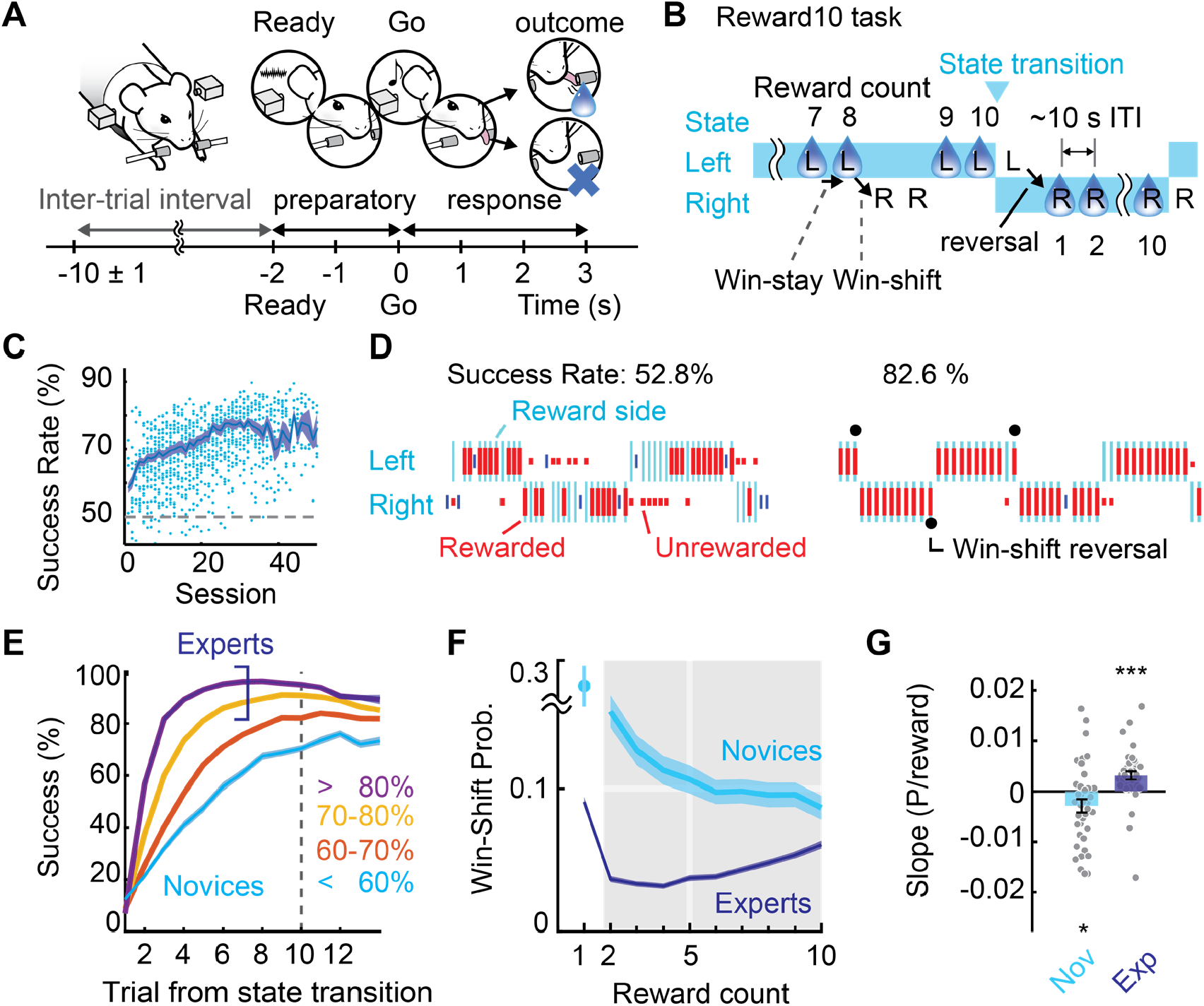
Mice learned to predict state transition. (**A**) Time-course of one trial. (**B**) State transition occurs after every 10^th^ reward delivery (Reward10 task). (**C**) Learning trajectory. (**D**) Example behaviors of low (left) and high (right) success rate animals. Vertical lines indicating states (cyan) and action-outcomes (long-red: rewarded, short-red: unrewarded, short-blue: premature lick, top/bottom rows: left/right choices). (**E**) Trial-by-trial success rate from the state transition in different learning stages. (**F**) Win-shift probability. (**G**) Slope of the win-shift probability in the shaded region in (F) (2nd to 10th reward counts) (novice, *t*_(37)_ = -2,16, *P* = 0.037; expert, *t*_(41)_ = 3.88, *P* = 3.7×10^−4^, *t-*test). Circles: individual animals. Mean ± SEM.

Mice reached the criterion performance (70% ≥ success rate, Fig. 1C) after a period of training. The behavior data was classified into two categories for comparison (novice, session success rate < 60%; expert, ≥ 70%; *n* = 103 and 675 sessions, respectively). Experts make reversals faster than novices (Fig. 1D and E; for premature rate and reaction time, fig. S1A and B). However, their trial-by-trial success rate peaked before the tenth trial in Reward10 task (fig. S1C, experts), indicating that experts did not select the previously rewarded action, rather selected an alternative (Fig. 1B; win-shift) instead of selecting the same action (win-stay). Indeed, the win-shift probability increased toward the state transition in experts (Fig. 1F and G) which occasionally led to reversals without mistakes (win-shift reversals, Fig. 1D filled circles). The increase of win-shift suggests that the animal predicted the approach of state transition. Further tests in five rewards per state task (Reward5 task) and short ITI task showed that mice make win-shifts much earlier in Reward5 task, and do not rely on the internal timing (Supplementary Text1, figs. S2 and S3). These results demonstrate that mice can predict the state transition triggered by the number of rewards delivered.

To understand the value dynamics behind this predictive choice, we resorted to the framework provided by reinforcement learning (*2*). Previous studies showed that the retrospective value can well explain the choice behavior in the dynamic foraging task (*3-5*). However, we found that retrospective value alone cannot account for the increase of win-shifts in our task (fig. S4). Instead, a model with action-value constructed from the hybrid of the retrospective and prospective values (Fig. 2A and fig. S5A, hybrid Q-learning [hQ-learning]; Methods) was sufficient to explain the increase of win-shift in experts (Fig. 2B, for model evaluation, fig. S5B and C). The contribution of prospective value was initially close to zero in novices but significantly increased in experts (fig. S5D and E, mixing weight *w*). This suggests that the novices rely on retrospective value, and experts incorporated the prospective value into their action selection. Our model predicts that the action-value increases monotonically as more reward is consumed in novices, but it develops into the rise-and-decay pattern in experts (Fig. 2C). This model also implies that the decay of action-value contributes to the increase of win-shifts in experts.

**Figure 2.**
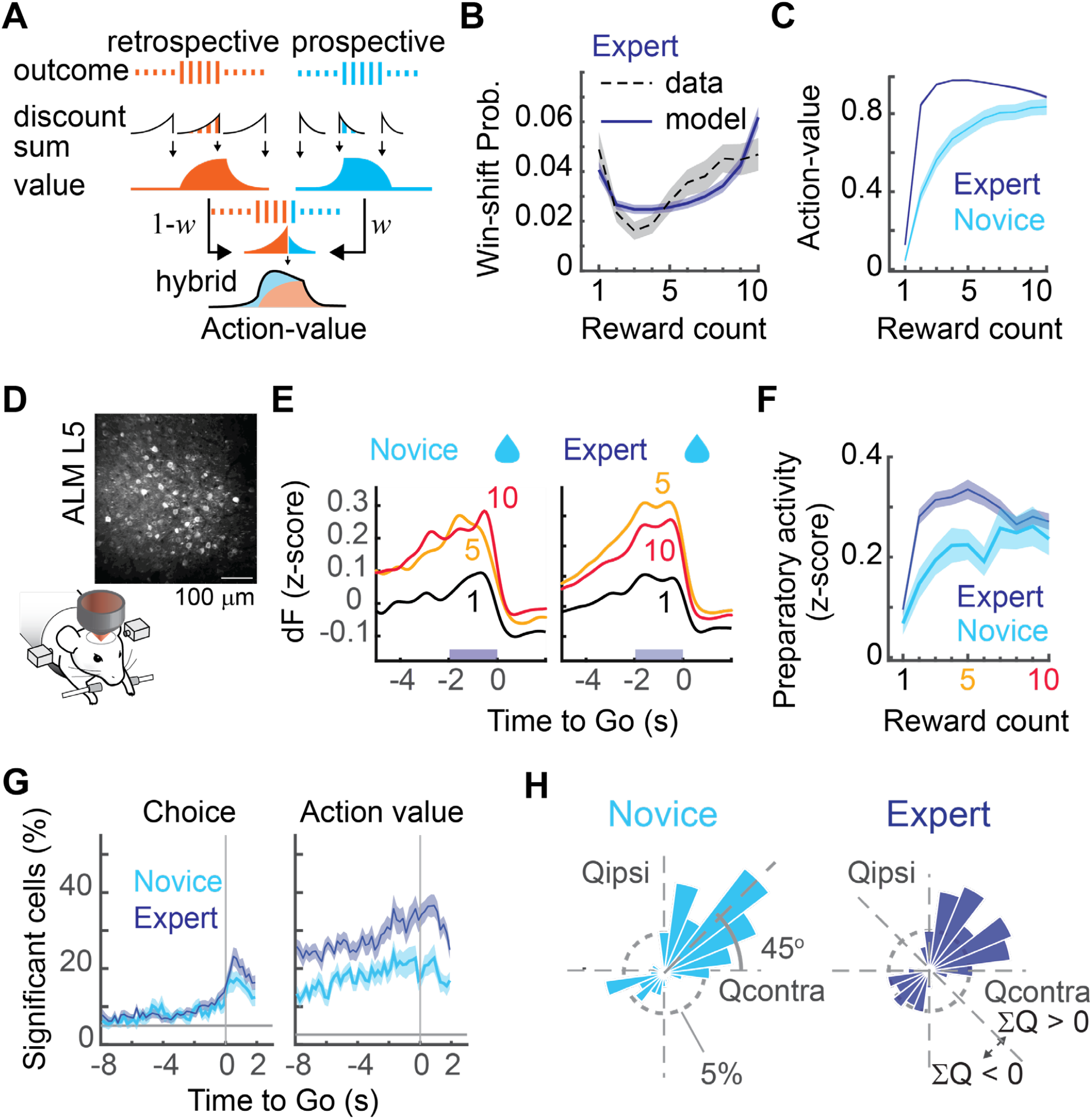
Emergence of prospective value in experts. (**A**) Hybrid Q-learning (hQ-learning) constructs action-value as a hybrid of retrospective and prospective values. (**B**) Win-shift probability of expert mice (dashed line) fit by hQ-learning (solid line). (**C**) Predicted dynamics of action-value. (**D**) Two-photon calcium imaging in ALM layer V (data selection criteria in table S1). (**E**) Trial-averaged activity (deconvolved fluorescence, dF) of ramping-up neurons in novices (Left, *n* = 156 cells from 21 image planes in 8 mice) and experts (Right, *n* = 459 cells from 30 image planes in 10 mice). Numbers are reward counts. (**F**) Modulation of preparatory activity (averaged over preparatory period) by reward counts. (**G**) Percentage of ramping-up cells significantly modulated by choice or action-value in novice and expert (multiple linear regression, *t*-test). (**H**) Polar bar plots showing the percentage of cells with a significant regression coefficient of action-values. Qcontra (Qipsi): action-value of the contra (ipsi)-lateral lick. Mean ± SEM.

To investigate whether the neuronal activities in ALM follow the model prediction, we conducted longitudinal two-photon calcium imaging in the ALM, targeting shallow layer V neurons (400-600 μm deep, Fig. 2D and movie S1; fig. S6 for imaging methods and single-cell examples). We sought the neural signature of the value dynamics in the ramping-up cells which showed the highest activity during the preparatory period (Ready to Go) and plotted the population average at various reward counts (Fig. 2E). This analysis revealed the clear concordance between the dynamics of preparatory activity and the predicted value dynamics in both novices and experts (Fig. 2C and F; for the unrewarded or unpreferred lick data, see fig. S7). A regression analysis further revealed that the majority of ALM ramping-up cells were action-value selective during the preparatory period (Fig. 2G (Action Value), novice, 20.7 ± 2.8 %, expert, 30.5 ± 2.7 %, *P*_threshold_ = 0.025; for history dependence, see fig. S8A and B) compared to imminent choice (Fig. 2G (Choice), novice, 9.1 ± 0.6 %, expert, 10.1 ± 0.6 %, *P*_threshold_ = 0.05; for non-ramping-up cells, fig. S8C). Among the action-value coding ramping-up cells, the majority was positively correlated to action-values in both novices and experts (ΣQ = Qcontra + Qipsi ≥ 0, novice 72.4%, *P* = 9.11×10^−10^; expert, 64.5%, *P* = 6.3×10^−10^, Binomial test. Fig. 2H and fig. S8H-M). A similar emergence of prospective value from retrospective one was also observed in the Reward5 task (Supplementary Text2). These results indicate that ALM preparatory activity initially encodes retrospective value, and prospective value emerges as the animal learns the state-transition structure of a decision-making task.

An implication of our model is that the decrease of action-value is the cause of win-shift choice. We then investigated the difference of action-value before win-stay and win-shift choice in experts. To minimize the effect of choice and reward history, we selected the same action-outcome sequence, uninterrupted 10-wins followed by either stay or shift (Fig. 3A, 10-wins & stay, and 10-wins & shift) and investigated the preparatory activity of positive action-value neurons (ΣQ>0) before each winning-action. We found that the action-value started to diminish several trials before changing the action but not when staying with the same action (Fig. 3B and C). A similar trend was also observed when the animal makes a predictive but premature reversal (9-wins & shift, Fig. 3D-F). The actions and outcomes are the same during the plotted range, therefore the faster decay of action-value is the precursor of the future win-shift choice, consistent with the model prediction.

**Figure 3:**
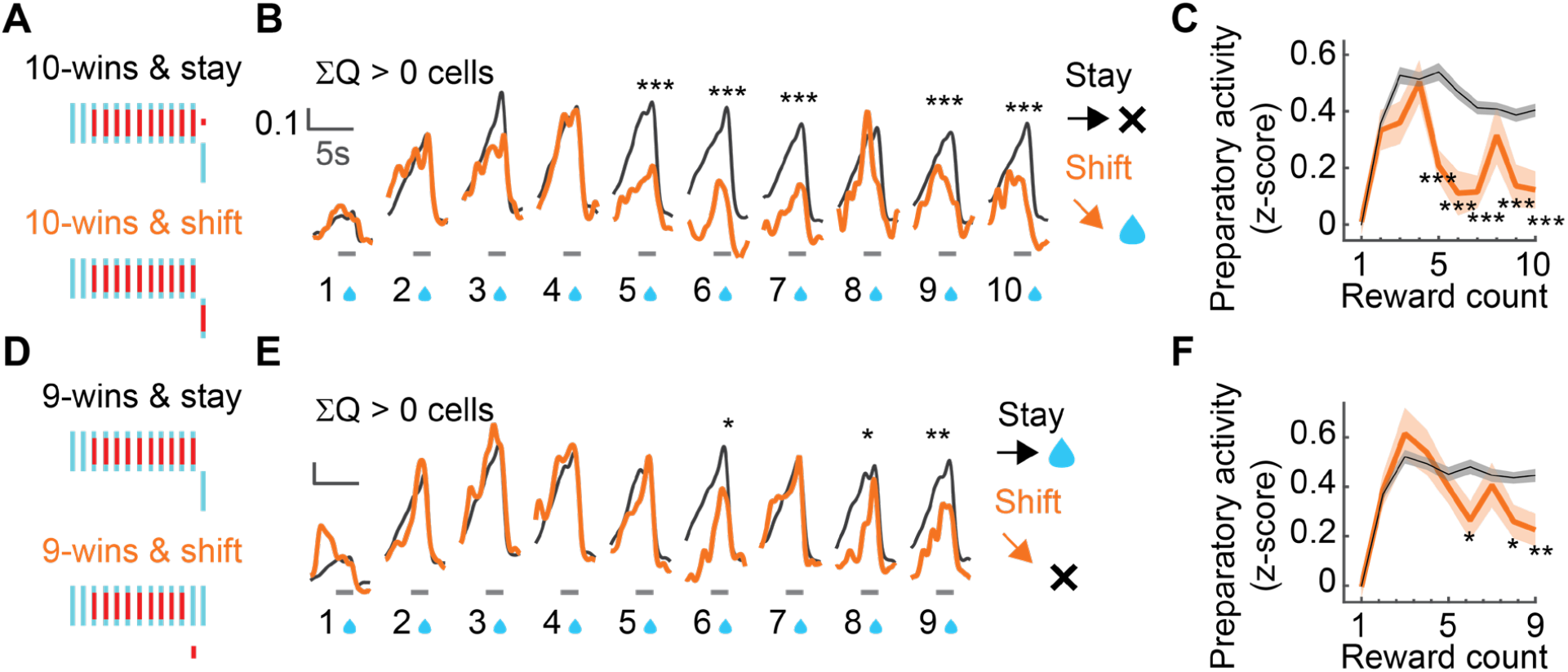
Value signal drops earlier when the animal makes predictive reversals. (**A**) Target choice sequences. (**B**) Preparatory activity during uninterrupted 10-wins followed by stay (10-wins & stay: black, *n* = 316 cells × trials) or shift (10-wins & shit: orange, *n* = 64). Gray bar, preparatory period. Data from positive action-value neurons (ΣQ>0) in experts. (**C**) Preparatory activity before each rewarding licks, *t*-test. (**D**,**E**,**F**) Same analyses for uninterrupted 9-wins & stay (*n*=316) and 9-wins & shift (*n*=73). *t-*test, \*\*\**P* < 0.001, \*\**P* < 0.01, \**P* < 0.05. Mean ± SEM.

Finally, to test the causal role of ALM preparatory activity on predictive choice behavior, we used photoinhibition of ALM (*18*) (Fig. 4A and fig. S9, *n* = 8 expert mice). ALM photoinhibition and control stimulation targeting the headpost were conducted in alternating blocks (every trial in 2 to 3 consecutive states for ALM photoinhibition; 3 to 7 consecutive states for control stimulation). We found that bilateral ALM inactivation during the preparatory period significantly delayed the reversal (Fig. 4B and C; fig. S10A) and suppressed the increase of win-shift probability (Fig. 4D and E) as if the experts were reverted to novices (compare Fig. 1E-G and Fig. 4C-E). The reaction time was not affected (fig. S10B). Similar photoinhibition during ITI (−10 to -5 s before Go) had no significant effect (fig. S10C). These results suggest that the preparatory activity in ALM plays a causal role in reflecting the prospective value into action selection.

**Figure 4:**
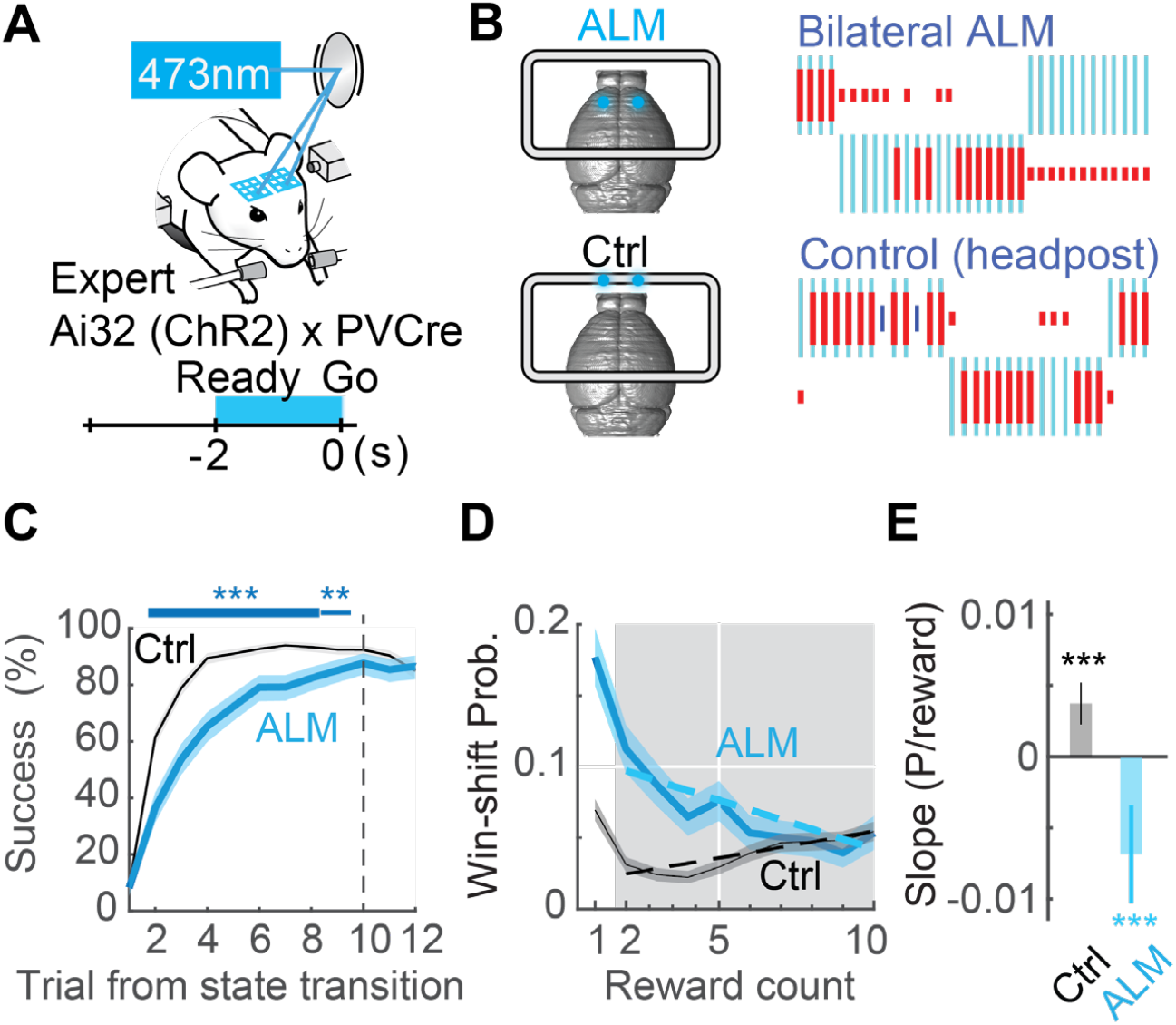
Photoinhibition of ALM preparatory activity inhibited predictive action selection in experts. (**A**) Schematic view of the photoinhibition experiment. (**B**) Example behaviors during the bilateral ALM photoinhibition and control blocks. (**C**) Trial-by-trial success rate from the state transition. Control (black) and ALM photo-inhibited (blue) trials. (Fisher’s exact test, \*\*\**P* < 0.001, \*\**P* < 0.01). (**D**) Win-shift probability in control (black) and ALM inhibited (blue) blocks. (**E)** Slope of win-shift probability (dashed line in (D), Ctrl: *t*_(9700)_ = 4.99, *P* = 5.98×10^−7^, ALM inhibition: *t*_(2980)_ = -3.89, *P* = 0.0001, linear regression, *t-*test). Mean ± SEM.

Previous studies mainly focused on the role of ALM in orofacial motor control (*18-20*). Although the value representation in ALM was reported (*5, 22*), they were not the main focus of their studies. In this study, we have identified ALM as the key cortical area that integrates the retrospective and prospective values. Photoinhibition of preparatory activity in experts selectively disturbed the structural-knowledge dependent behavior (e.g., increase of win-shifts), sparing the novice-like behavior (e.g., slow reversal) likely to be maintained outside of ALM, such as the regions in the posterior cortex that involves in history-dependent behavior (*5, 16*). Together, our results show that the frontal cortex (ALM) is the critical hub to inject structural knowledge into action selection and to promote predictive decision making.

## Acknowledgments

We thank Mitsuko Uchida, Naoshige Uchida, Kenji Doya, James Hejna, and Richard Mooney for their comments. Bernd Kuhn for his advice on two-photon microscope setup. Satoshi Yawata constructed the pAAV-CaMKIIa-GCaMP6s plasmid. Fumi Ageta assisted with animal training. pAAV-CaMKIIa-GCaMP6s-P2A-nls-dTomato was a gift from Jonathan Ting (Addgene plasmid # 51086).

## Funding

MEXT/JPSP KAKENHI 19H04983 and 21H02804 (KH)

MEXT/JPSP KAKENHI JP18H04014 (DW)

JST CREST JPMJCR1756 (DW)

Takeda Science Foundation (DW)

## Author contributions

Conceptualization, Methodology, Visualization, Formal Analysis: KH

Investigation: KH HTA

Writing – Original Draft: KH

Funding Acquisition Resources: KH DW

Supervision: DW

## Data and materials availability

HDBCellSCAN is deposited to GitHub (https://github.com/hamaguchikosuke/HDBCellSCAN). Imaging and electrophysiological data will be shared upon a reasonable request.

## Supplementary Materials

Materials and Methods Supplementary Text1 to Text2 Figs. S1 to S10

Tables S1

References (*1-23*), (*24-29)*

Movies S1 to S2

## Supplementary Materials for

## Materials and Methods

### Animals

All experiments were approved by the Animal Research Committee, Graduate School of Medicine, Kyoto University. C57BL/6N mice (*n* = 35, Japan SLC) and PV-Cre × Ai32 mice (*n* = 8, PV-Cre: B6;129P2-*Pvalb*^tm1(cre)Arbr^/J [JAX 008069]; Ai32: B6.Cg-*Gt(ROSA)26Sor*^tm32(CAG-COP4*H134R/EYFP)Hze^/J [JAX 024109]) were used. All mouse strains used in the behavioral studies were maintained in a C57BL/6N background. Mice were individually housed in an aluminum cage with an environment enrichment apparatus (a running wheel). The room was in a reversed light cycle (12h - 12h) and all the experiments were conducted in the dark phase. Male mice aged 8 weeks or older were used in all experiments.

### AAV preparation

The plasmid carrying the CaMKIIa promoter and GCaMP6s sequence (pAAV-CaMKIIa-GCaMP6s) was derived from pAAV-CaMKIIa-GCaMP6s-P2A-nls-dTomato (addgene plasmid #51086) by deleting the P2A-nls-dTomato sequence. Recombinant adeno-associated virus (AAV) expressing GCaMP6s under the control of the CaMKIIa promoter was packaged in serotype AAV2/9. AAV was purified by iodixanol gradient ultracentrifugation. The final concentration was assessed using realtime PCR (original titer 1.2×10^13^ vg/ml).

### Surgery

Headposting surgeries were conducted at least three weeks before the behavior training. Mice were anesthetized with isoflurane (1-2%) and placed in a stereotaxic device, followed by subcutaneous injection of dexamethasone (2 mg/kg). Body temperature was maintained using an electric heater. The scalp was dissected out in a round shape, and the periosteum over the dorsal surface of the skull was removed. The imaging areas were identified according to the stereotaxic coordinates. The target was marked on the surface of the skull with a razor blade and painted with a black marker. A rectangular stainless-steel frame (CF-10, Narishige) was attached with clear dental cement (Superbond C&B, Sun Medical). A single craniotomy was made over both cerebral hemispheres. A glass pipette attached to a Nanoject-II (Drummond Scientific) was used to deliver AAV expressing GCaMP6s (AAV-CaMKII-GCaMP6s, diluted to 2.4×10^12^ vg/ml, 40-60 nl per imaging site). A glass window (5mm diameter, No. 2 thickness, Matsunami-Glass) was cut to fit the craniotomy and was secured on the remaining edge of the skull by using the same clear dental cement. After the surgery, Enrofloxacin (2.5% Baytril, Bayer, diluted to 8 μl/ml) and Carprofen (Rimadyl, Zoetis JP, diluted to 1.6 mg/ml) were administered via drinking water for at least 5 days. After that, mice were allowed to recover with free access to water for two weeks before water restriction. A subset of mice received windowing surgery after they reached the criterion (70% success rate), followed by at least 3 days of a recovery period. For photo-inhibition experiments, the same procedure was used except for the craniotomy. The skull was only covered with the same clear dental cement. For electrophysiological recording experiments, several days before the recording, a craniotomy (∼1 mm diameter) was made over the skull, and a ground and a reference pin were implanted over the cerebellum. The surface was covered by silicon elastomer (Kwik-Sil, WPI) to protect the craniotomy until the recording. The extracellular activity was recorded using a silicon probe (A1×32 series, NeuroNexus Inc) through a head-stage amplifier (RHD-2000, Intan). Photo-stimulation patterns and electrophysiological data were simultaneously recorded via a data acquisition board (RHD USB Interface Board, Intan) and analyzed by custom-written software (by KH, MATLAB, Mathworks).

### Pre-training

Mice received pre-training before performing a serial reversal-learning task. Two to three days before the start of pre-training, mice were water restricted at 1 ml per day and acclimated to the handling of an experimenter. The preferred side of licking was assessed by manual water feed. A behavior control program running in real-time (written in MATLAB and Simulink Realtime by KH) was used to control and record behavioral events. Initially, the rewarding side was fixed to the unfavored side. The head-fixation period started at less than 5 min, and eventually extended up to 90 minutes over the sessions. When the mice reached the first criterion (more than 100 trials within 30 min), licks between “Ready” and “Go” signals were categorized as premature licks: they were punished by a loud noise and the state of the trial reverted to the inter-trial interval. Once the percentage of premature trials became less than 15% of all the trials, the water delivery port was changed to the other, originally favored side. This condition was kept fixed until the same criterion (the percentage of premature trials became less than 15% of the total trials) was met. This phase of pre-training usually lasted for 5-20 days.

### Deterministic State Transition Task

Once the mice learned to lick both sides of the water ports with minimum premature behavior as described above, we initiated the deterministic state transition task. In the main task, the mice were required to refrain from licking after hearing the “Ready” noise and to wait until the “Go” tone presentation. Any lick between “Ready” and “Go” signals was punished by the loud noise, and the trial was considered as a premature one. Both novice and expert mice performed the task with a small fraction of premature actions with a similar level of reaction time (fig. S1A and B). After the “Go” tone presentation, the first detected lick within the response window (3 s) was considered as the response. The detection of response was reported to the animal as a brief presentation of a click sound (∼100 ms). In the Left (Right) state, the left (right) response was considered as a “correct” response and rewarded with a drop of water (∼4 μl). On the other hand, in the Left (Right) state, the right (left) response was considered as an “incorrect” response, and no water was supplied. The rewarding water port was changed when the mice gained a prefixed amount of reward (10 for the Reward10 task, 5 for the Reward5 task). Reward delivery is deterministic and was always given for the correct responses, and no reward was given for the incorrect responses. No lick within the response window was considered as a missed trial, and water was not supplied.

Initially, the mice were trained with the 10 rewards per state condition (Reward10). A subset of mice was also subjected to a 5 rewards per state (Reward5 task) task if they had already become Reward10 experts (at least 70% success rate) over 3 consecutive days in the past. One session lasted maximally 90 min or until the mice obtained 1.5 ml of water. If mice did not maintain stable body weight, they received supplemental water. The main training lasted 50 to 190 sessions. We did not observe any decline of fluorescence of genetically encoded calcium indicators (GCaMP6s) during this period.

### Behavior Control Hardware and Software

The behavior experiments under two-photon microscope were performed in a dual-lick operant module (OPR-1410, O’Hara & Co., LTD). The rest of the behavior training and optogenetic experiments were performed using in-house built dual-lick operant chambers. The onset of the lick was detected by the break of infra-red beam situated in front of the water delivery port. The auditory stimuli were presented by a pair of miniature speakers (FK-23451-000, Knowles) attached to the head-fixation stage (O’Hara & Co., LTD) near the ears of the mouse. Water delivery was controlled by a solenoid valve (LHDA1233315H, The Lee Company). The duration of the valve opening was adjusted to deliver 4 μl of water. The lick ports and mouse chair were designed with a 3D CAD software (Solidworks, Dassault Systems) and printed in a 3D printer (Form2, Formlabs).

In all the experiments, a custom-written behavior control program running in real-time (written in MATLAB and Simulink Realtime by KH) was used to control the behavioral events. The behavioral events, imaging scan signal, and laser control signals were recorded at 1kHz by using a data acquisition board (MF634, Humusoft or PCI-6251, National Instruments) and streamed to the host PC.

### Behavior Data Analysis

The session success rate and the trial-by-trial success rate were defined as the fraction of rewarded trials, excluding the misses and premature trials. The error rate was defined as 1 – success rate. The win-shift probability was defined as the fraction of the win-shift actions among all wins (rewarded trials). The slope of win-shift probability was estimated from the pooled trials of each animal in Fig. 1, or pooled trials from all the optogenetically manipulated animals in Fig.4, using linear regression. The peaks of the trial-by-trial success rate were calculated for each session, and mean, median, quartiles were computed over the sessions. The reaction time was defined as the time from the Go signal to the first lick.

### Two-photon calcium imaging

Imaging experiments started after the mice had reached the criteria in the pre-training stage. Imaging was conducted using a commercial two-photon microscope (FVMPE-RS, Olympus) with a 25× objective lens (XLPN25XWMP2, Olympus). The beam of a femtosecond pulsed laser with a peak wavelength at 920 nm (Chameleon Vision II, Coherent) was delivered via a custom-built light path. The laser intensity was controlled by an acousto-optical modulator (AOM; MT110-B50A1,5-IRHk, OEM ver. AA Optoelectronic). Images (512 × 512 pixels covering a 500 × 500 μm area) were continuously recorded, up to 64000 frames at 30 frames per second, using image acquisition software (FV315-SW, Olympus). The average excitation power was up to 120 mW for deep layer imaging (∼ 500 μm deep).

### Pre-processing for Ca imaging data analysis

Data analysis was performed in MATLAB (Mathworks) and Python 3.6. Brain motion was detected as the peak of phase correlation between the mean image and each image. The detected motion was corrected by shifting each image in an x-y coordinate (no shear, no rotation) by using an algorithm in Suite2P (version 2016)(*24*). Regions of interest (ROIs) corresponding to the cell bodies were detected by using a custom-written algorithm (HDBCellSCAN). The fluorescence signal was deconvolved using a non-negative deconvolution algorithm (*25*) and was defined as the deconvolved fluorescence signal (dF in the main text).

### Ca imaging data analysis

All data analysis was performed using custom-written codes in MATLAB (Mathworks). First, neurons were tested for their selectivity to four trial types (Contra/Ipsi-lick × Rewarded/Unrewarded) during the preparatory period (−2 to 0 s before the Go signal), action period (0 to 1 s after Go), and outcome period (1 to 4 s after Go), using the deconvolved fluorescence (dF) averaged within each period (ANOVA, *P* threshold = 0.05). The ramping-up cells were selected as the cells that showed maximum activity during the preparatory period in any of the four trial types. For the rewarded trials, the data were grouped by the learning stage (Novice, Expert) and the cumulative number of rewards (reward count) within the state, and then averaged within the group. Note that the first reward trial was defined as the trial in which the animal receives the first reward within the state. For the unrewarded trials, the data were grouped by the learning stage and the cumulative number of unrewarded trials within the state and then averaged. In all the analyses above, the preferred lick direction of a neuron was defined as the direction in which the mean firing rate of rewarded trials was higher.

### Photoinhibition

Light from a 473 nm laser (OBIS 473 LX 75 mW, Coherent) was first introduced to an f = 100 mm lens (AC254-100AB, Thorlabs) to form a focused spot on a two-axis scanning mirror (Integrated MEMS Mirror, Mirrorcle Tech), and was subsequently re-focused onto the skull surface by another f = 75 mm lens (AC254-75AB, Thorlabs). The laser had a Gaussian profile with σ = 160 μm at the level of the skull surface. The laser spot size on a piece of black paper was measured by a CMOS camera (DMK37BUX287, ImagingSource).

The MEMS mirror directed the light in a step-wise manner to stimulate multiple brain regions sequentially. The MEMS mirror was aimed at each target for 50 ms. Transient time was less than 2 ms. The laser intensity was controlled through a remote controller (OBIS Single Remote, Coherent) in the analog modulation mode. During the laser stimulation, the amplitude of the laser was sinusoidally modulated at 40 Hz and was linearly attenuated for the last 100-200 ms period. During the transient time of the mirror, the laser beam was transiently turned off to prevent stimulating the non-target area. Laser power was calibrated using a laser power meter (Fieldmate, Coherent). The laser intensity was set to 1.5 mW per spot throughout the experiments. All the photo-stimulation was done through clear dental cement and an intact skull. The attenuation level through the skull was 28.4 ± 4.8 % (*n* =5 skulls, mean ± SEM) measured by using the power meter and the skull after the experiments. This corresponds to 0.42 ± 0.07 mW per spot in our experiments.

Photoinhibition experiments were conducted in alternating blocks; one block corresponds to 2 to 3 consecutive states for ALM bilateral photoinhibition and 3 to 7 consecutive states for control stimulation. In ALM targeting blocks, the laser was aimed at two spots over ALM (1.5 mm left and right from the midline, 2.5 mm anterior from the bregma). In the control blocks, the laser was aimed at the frontal edge of the head post (two spots at 1.5mm left and right from the midline, usually 3.5 to 4.5mm anterior from bregma). The length of states within one block was randomly chosen. The experiments were always started with a control stimulation block.

### Electrophysiological data analysis

Data analysis was performed in MATLAB (Mathworks) and Python 3.6. First, isolated units were identified from the multi-channel recording data by using Kilosorts (*26*) and manually re-clustered by using Phy (https://github.com/cortex-lab/phy). The difference in the firing rate during photo-inhibition and control periods was used to assess the range of photo-inhibition effects.

### Statistical Analysis

All the statistics were computed using MATLAB (Mathworks). All test for differences were two-sided and differences were considered as significant when *P* < 0.05. Bonferroni correction was applied as necessary.

### Reinforcement Learning Models

#### Retrospective Value by Q-learning

The retrospective value, the discounted sum of past rewards obtained by an action *a* after the observation of outcome at trial *i* is given as follows:

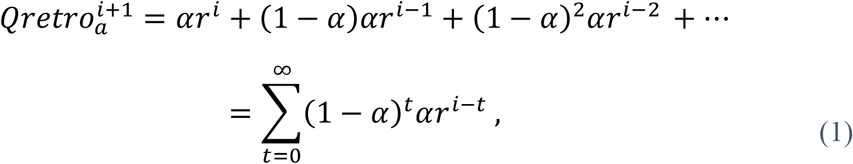

where 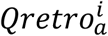 is the retrospective value of the chosen action *a* ∈ {*Left,Right*} at the *i* -th trial, 1 − *α* (0 ≤ *α* ≤ 1) is the discounting factor for past rewards. The outcome at trial *i* is *r*^*i*^ ∈ {0,1}, (1 for rewarded chosen action, and 0 otherwise). Retrospective value *Qretro* is then limited within [0 1] range. Caching the infinite history of reward is impractical, but one could rewrite Eq. (1) to update the retrospective value as follows;

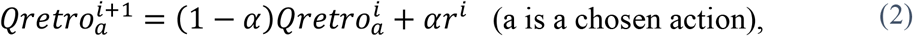

which depends only on the previous value and the latest outcome. This update rule is equivalent to Q-learning, a standard model-free reinforcement learning. The modified version of Q-learning that includes the forgetting term for unchosen action,

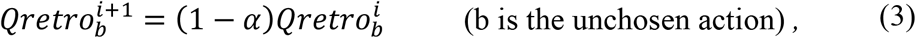

and choice bias term was used to explain rodents’ choice behavior in the dynamic foraging task (*10, 27*). The action selection probability is assumed to be the soft-max function, and for the binary choice case, it simplifies to

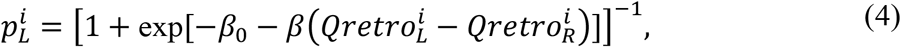

where *p*_*L*_ represents the probability of choosing left, and the probability of choosing right is 1 − *p*_*L*_. The inverse temperature *β* is a positive scalar that controls the randomness of the choice. The choice bias term is represented by *β*_0_. This model provides the retrospective value in this study.

#### Prospective value in hybrid Q-learning

To incorporate the knowledge about the task, we considered a hybrid model that incorporates both the discounted sum of past and future rewards. In the deterministic state transition task, the “true” value (the upper limit of the discounted sum of future rewards) of the currently rewarding state can be analytically calculated as follows:\

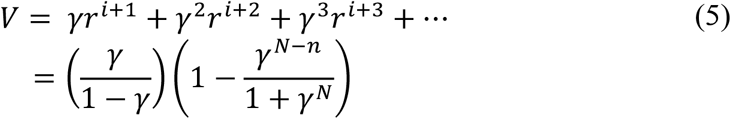

where γ (0 ≤ *γ* ≤ 1) is the discount factor of future reward, *N* is the maximum number of rewards per state (10 for Reward10, and 5 for Reward5 task), *n* is the number of rewards obtained in the current state. We will then construct the hybrid value as the mixture of retrospective and prospective value. We normalized *V* to take [0 1] range, and defined the prospective value *Qpros* as below:

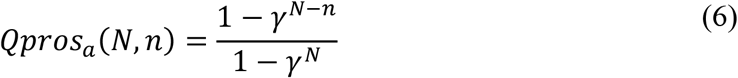

The prospective value for unrewarding action is 1 − *Qpros*_*a*_(*N, n)*. The hybrid value (fig. S5A) is defined as

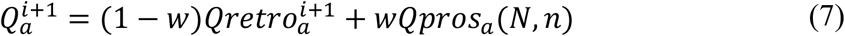

where w (0 ≤ w ≤ 1) is the mixing weight. When *w =* 0, hybrid Q-learning is reduced to Q-learning. The action selection mechanism is the same as Q-learning, represented by the softmax function as below:

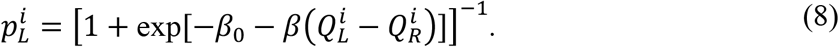

#### Model Evidence

We fit the whole trajectory of learning by a single learning model while allowing the parameters to evolve over sessions. To select a better model while avoiding the problem of overfitting, various approximated criteria, such as Akaike Information Criterion (AIC), Bayesian Information Criterion (BIC), were proposed. BIC can be derived as an approximation of the model evidence, the likelihood of a statistical model marginalized by the prior distribution of model parameters. Model evidence is by construct not biased to the given data as the maximum likelihood estimate (MLE) does. The ratio of the model evidence is called the Bayesian factor and is used as a criterion in the model selection (*28*). Recent advancements in the computational sampling method allow direct computation of the model evidence. Here we fit each session data by Markov Chain Monte Carlo (MCMC) method with Metropolis-Hasting algorithm (*29*). This procedure directly produces the sample distribution of model parameters. Then we can obtain the model evidence (marginalized likelihood) as the sample average of MCMC samples:

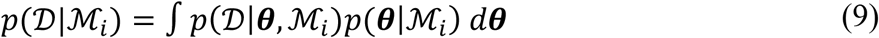

where 𝒟 is the data (choice and reward sequence in one session), ℳ_*i*_ is the label of i-th model, ***θ*** is a vector of free parameters of the model ({*α, β, β*_0_} for Q-learning, and {*α, β, β*_0_, *γ, w*} for hybrid Q-learning), *p*(𝒟 |***θ***, ℳ_*i*_) is the likelihood of the model, and *p*(***θ***|ℳ_*i*_) is the prior distribution of the parameters. For the initial session data, the prior distribution is set to *beta*(2,2), log-normal(2,1), normal(0,1), *beta*(2,2), *beta*(1,2) for {*α, β, β*_0_, *γ, w*}, respectively. Here, *beta*(a,b) is the beta-distribution (not the inverse temperature). After the first session, the prior distribution of the model parameter at *k+1* th session is given as

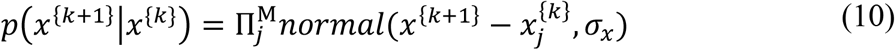

where 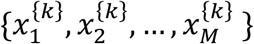 is a set of MCMC samples (M = 1000) from the *k*-th session, *normal*(*μ, σ*) is the normal (Gaussian) distribution, *σ*_*x*_ is the hyper parameter that controls the spread of parameters between two sessions and is set to {0.1, 0.3, 0.3, 0.1, 0.1} for {*α, β, β*_0_, *γ, w*}, respectively. The convergence of distribution was tested by the following procedure: After 30 initial steps, MCMC samples were divided into 8 clusters using k-means clustering. Then ANOVA was used to test the null hypothesis that there is no difference in the mean of all the clusters. The convergence was defined as the state where all the model parameters have p-values above p > 0.05.

### Regression Analysis

We performed a multivariate regression analysis of the deconvolved fluorescence activity to test whether the neuronal activity was correlated with any of the behavior events and latent variables of the model. The regression model was

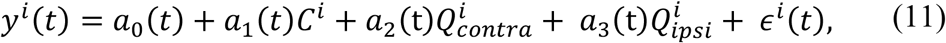

where *y*^*i*^ (*t*) is the deconvolved fluorescence during the *i*-th trial, *C*^*i*^ is the choice during the *i*-th trial (contra- or ipsilateral choice; 1 or -1), 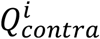 and 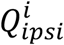 are action-values associated with the contra- and ipsilateral actions before observing the outcome of the *i*-th trial. The residual error term is *ϵ*^*i*^ (*t*), and *a*_0_, … *a*_3_ are the regression coefficients. The median variance inflation factors for choice, 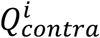 and 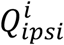 were 1.62, 1.65, and 1.45 in novice, 3.32, 2.61, and 2.32 in experts.

We also used the following model to regress the contribution from the past choices and outcomes in the deconvolved fluorescence activity;

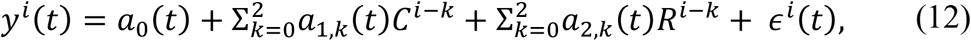

where *R*^*i* −*k*^ represents rewarded (1) or unrewarded (0) on the (*i-k*) th trial, and *a*_*n,k*_ represents the regression coefficients of past choices and outcomes. The median variance inflation factors for the choice at current trial (*a*_1,0_), one trial before (*a*_1,1_), two trials before (*a*_1,2_), the reward outcomes at current trial (*a*_2,0_), one trial before (*a*_2,1_), and two trials before (*a*_2,2_), were 2.10, 2.58, 2.02, 1.86, 2.22, 1.80 in novice, and 2.44, 3.63, 2.42, 1.29, 1.59, 1.35 in experts.

### Histology

Mice were perfused transcardially with PBS followed by 4% PFA/0.1 M PB. The brains were postfixed overnight and transferred to 30% sucrose until sectioning on a microtome (Leica, CM1850). Coronal, 40 μm free-floating sections were collected in PBS-azide. Slide-mounted sections were imaged on a Keyence microscope (BZ-X700).

For the identification of PV-positive and ChR2-EYFP positive neurons, we used Rabbit-anti-GFP (Invitrogen A11122) and Mouse-anti-PV (Swant PV235) as primary antibodies (1:1000 dilution), and Goat-anti-Rabbit Alexa488 (Invitrogen), Goat-anti-Mouse Alexa 594 (Invitrogen) secondary antibodies (1:200 dilution). DAPI was used for counterstaining. For cell counting, we manually selected cells using the ImageJ multipoint selection tool (NIH).

## Supplemental Figures and Tables

**Figure S1.**
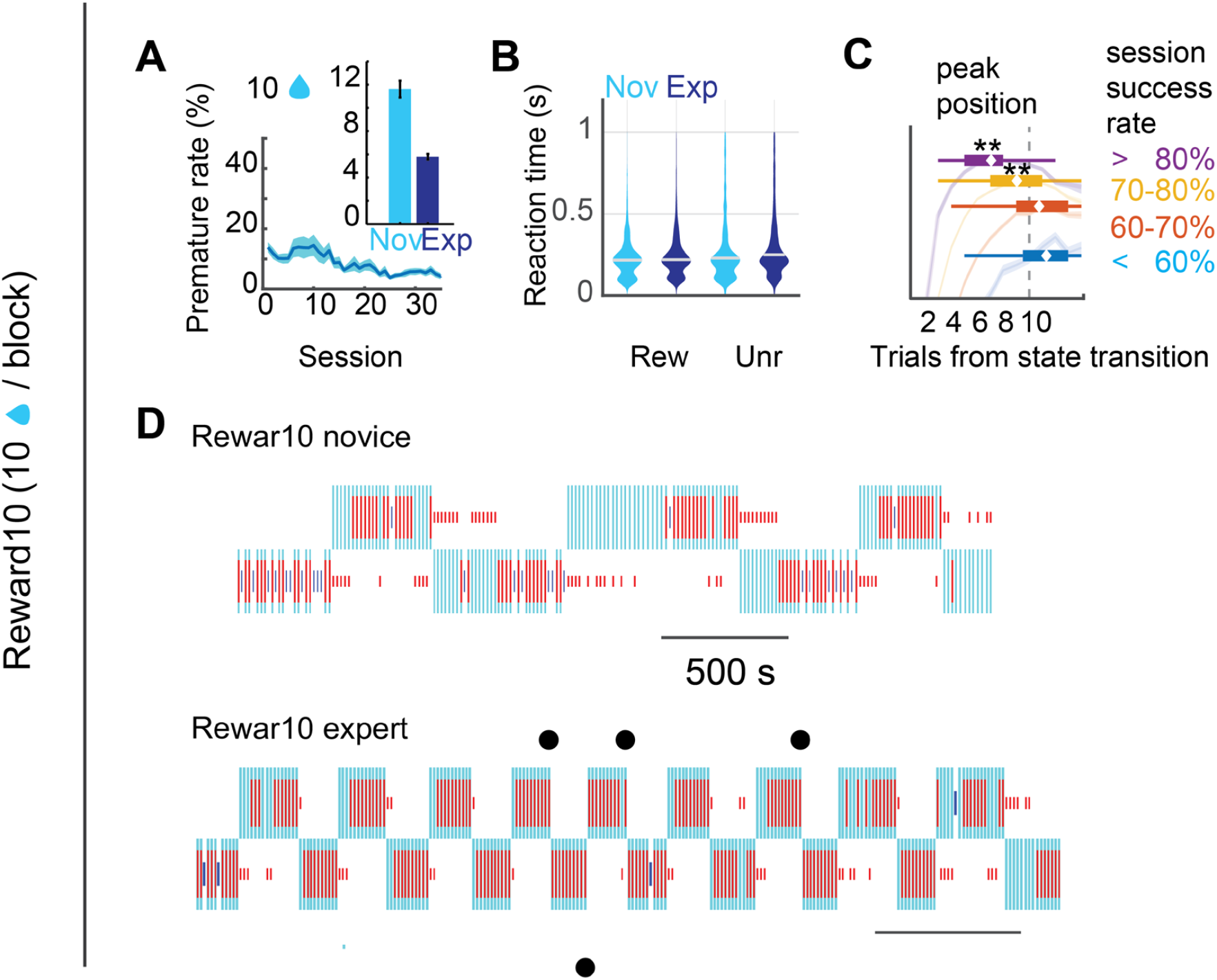
Reward10 behaviors. (**A**) Percentage of premature trials were both minimized in novices and experts. Inset: mean premature rates in each training stage. (**B**) Distribution of reaction time (Go cue to the first lick) before each rewarded (Rew) and unrewarded (Unr) action were highly similar in novices and experts; Novice (Rew), 212 ± 1.1 ms, *n* = 21420 actions; Expert (Rew), 217 ± 0.5 ms, *n* = 127260; Novice (Unr), 237 ± 5.9 ms, *n* = 22934; Expert (Unr), 248 ± 3.5 ms, *n* = 56898. Median ± SEM. (**C**) Peaks of the success rate were significantly earlier than 10 trials in Reward10 task in animals with 70-80% success rate (*P* = 2.6×10^−13^, Mann-Whitney *U*-test) and above 80% success rate (*P* =1.0×10^−6^). (**D**) Example of novice (top) and expert (bottom) behavior in Reward10 task. Red bar: long, rewarded; short, unrewarded. Cyan: reward side, not revealed to the animal. Top/bottom rows: left/right licks. Blue short bar: premature trial. Filled circles: win-shifts at the state transition.

**Figure S2.**
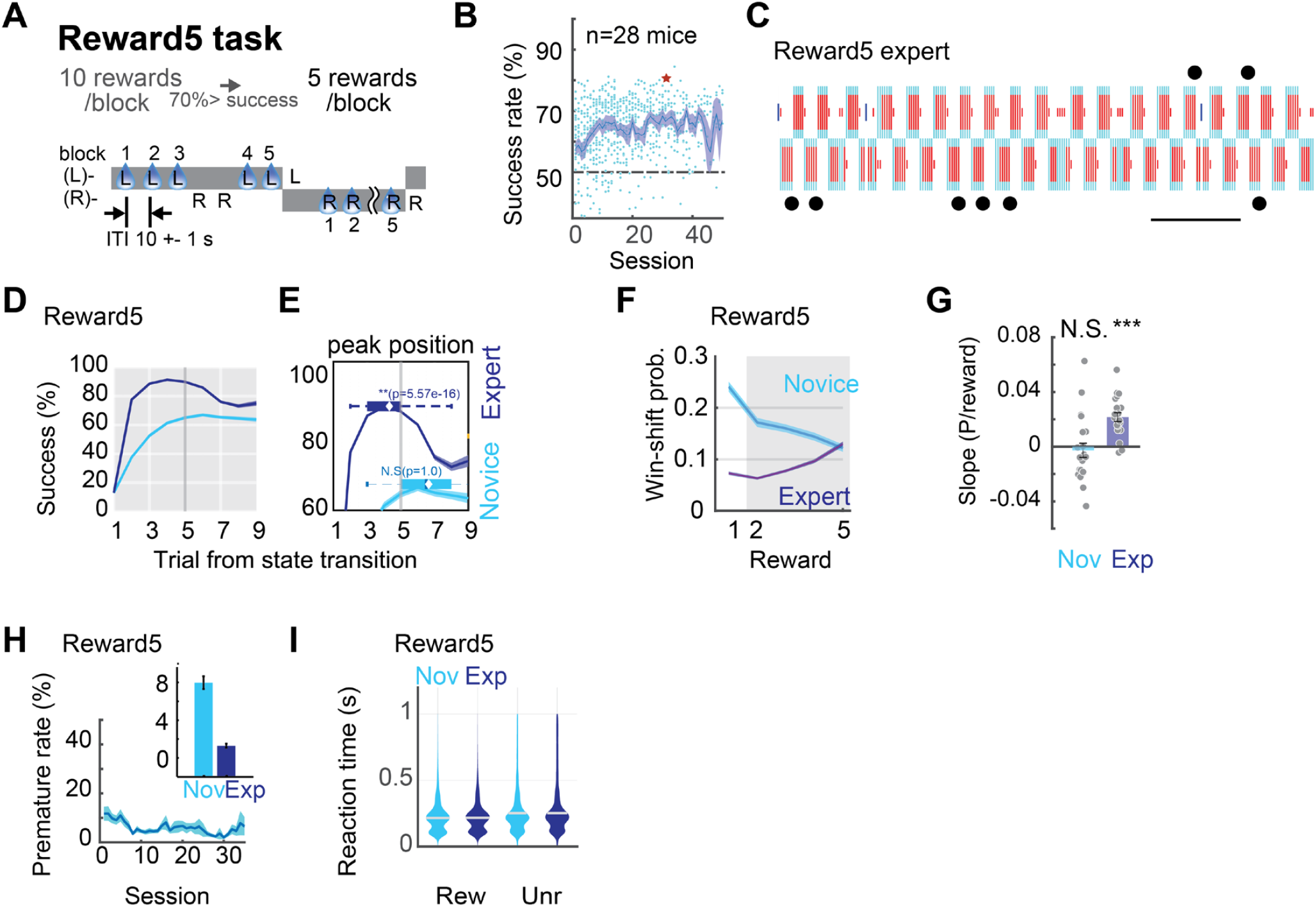
Reward5 task summary. (**A**) Reward5 task. The state (reward condition) transition occurs every five rewards. Reward10 experts were introduced to the Reward5 task. (**B**) Learning trajectory in Reward5 task. (**C**) An example of Reward5 expert behavior (star in (B)). Circles are win-shift reversals without mistakes. (**D**) Trial-by-trial success rate from the state transition in different learning stages. (**E**) Box plots representing the mean (diamonds) and first or third quartile points of the peak of the success rate calculated from each session. Peaks of the success rate were significantly earlier than 5 trials in Reward5 task in animals in experts (*n* = 260 sessions, *P* = 5.57×10^−16^, Mann-Whitney *U*-test) but not in novices (*n* = 180 sessions, *P* = 1.0). (**F**) Win-shift probability in different learning stages. (**G**) Slope analysis calculated from the shaded region in (F). Win-shift probability significantly increased from novice to expert (novice, *t*_(21)_ = -0.52, *P* = 0.68, *n* = 22 mice; expert, *t*_(20)_ = 6.77, *P* = 1.38×10^−6^, *n* = 21 mice, linear regression, *t*-test). (**H**) Rate of premature trials in Reward5 task. (**I**) Distribution of reaction time before each rewarded (Rew) and unrewarded (Unr) action in Reward5 task. Novice (Rew), 216 ± 1.1 ms, *n* = 22470 actions; Expert (Rew), 218 ± 1.0 ms, *n* = 28300; Novice (Unr), 253 ± 7.3 ms, *n* = 16026; Expert (Unr), 253 ± 6.9 ms, *n* = 19192. Median ± SEM.

**Figure S3.**
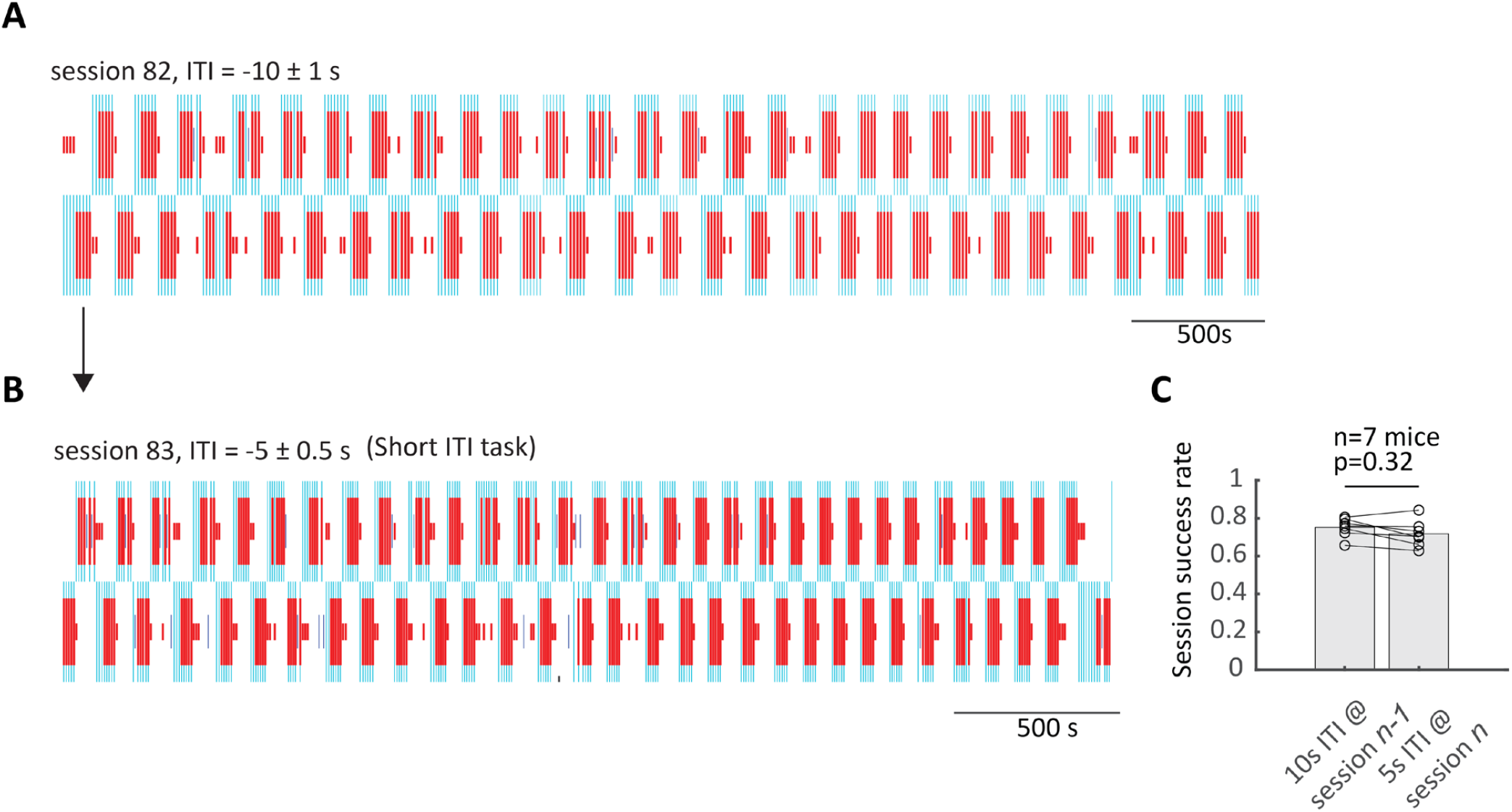
Short inter-trial interval (ITI) task. (**A**) An example choice behavior in the normal ITI (10 ± 1 s) task one session before the challenge day. (**B**) An example choice behavior in a short ITI (5 ± 0.5 s) task. (**C**) Session success rate of normal and short ITI tasks. No significant difference was detected. Paired *t*-test, *n* = 7 mice (*P* = 0.32).

**Figure S4.**
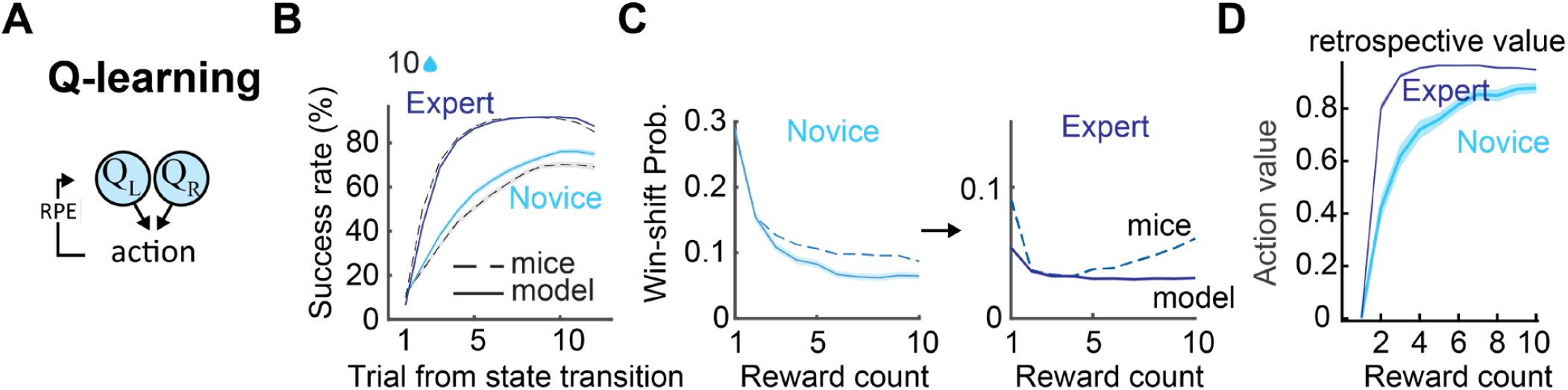
Retrospective value cannot explain win-shift behavior. (**A**) Schematic view of Q-learning. RPE: reward prediction error. For more details, see Materials and Methods. (**B**) Trial-by-trial success rate of the mice (dashed line) and simulated behavior (solid line) of the best-fit Q-learning aligned to the state transition. Expert (blue) and novice (cyan). (**C**) Simulated and experimental win-shift probability as a function of reward counts within a state. Note that in experts, model (solid line) cannot recreate the increase of win-shifts. (**D**) Predicted dynamics of Q-learning action-value (retrospective value) as a function of reward counts within a state.

**Figure S5.**
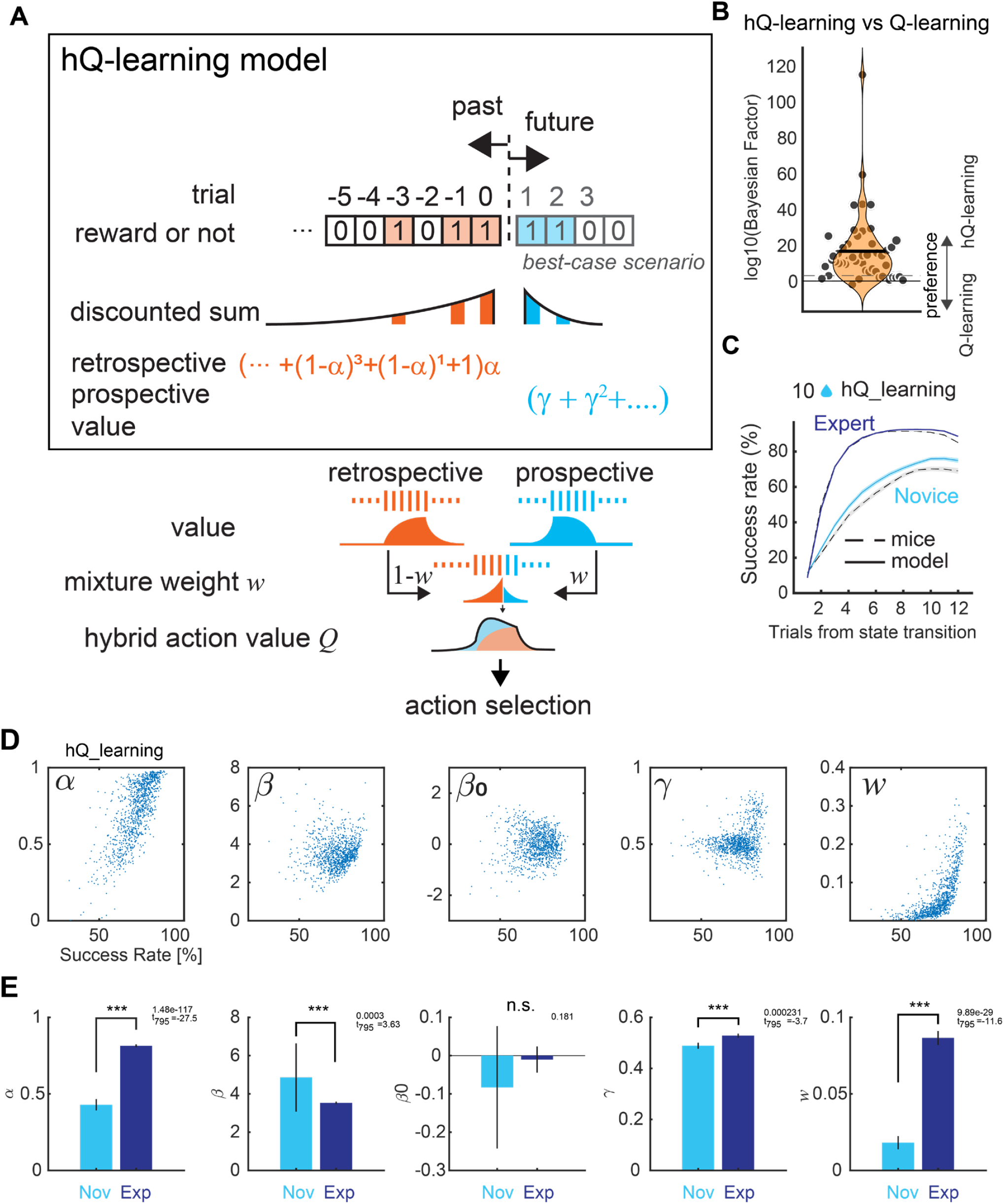
Hybrid Q-learning (hQ-learning) shows a better fit to the behavior of animals. (**A**) Schematic of hQ-learning model, showing the process to compute (hybrid) action-value for an action (e.g., left lick). See **Reinforcement Learning Models** in Methods for mathematical formulations. (**B**) Distribution of the Bayesian Factor showed that the hybrid of retrospective and prospective value (hQ-learning) explains the choice behavior of the mice better than the retrospective value alone (Q-learning). log10(BayesianFactor) = 16.6 ± 2.8, *n*= 43 mice. Dashed horizontal line indicates log10(BayesianFactor)=3 level where the posterior probability of data being generated from hQ-learning is 1000 times higher than Q-learning. Each dots represents a single animal. (**C**) Trial-by-trial success rate of the mice (dashed line) and simulated behavior (solid line) of the best-fit hQ-learning aligned to the state transition. Expert (blue) and novice (cyan). (**D**) Maximum a posteriori (MAP) estimates of meta-parameters for each session in Reward10 task. (**E**) Meta-parameters in different learning stages. The increase of prospective weight *w* (the rightmost panel) indicates that the animals incorporate the prospective value into their choices. Data are mean ± SEM.

**Figure S6.**
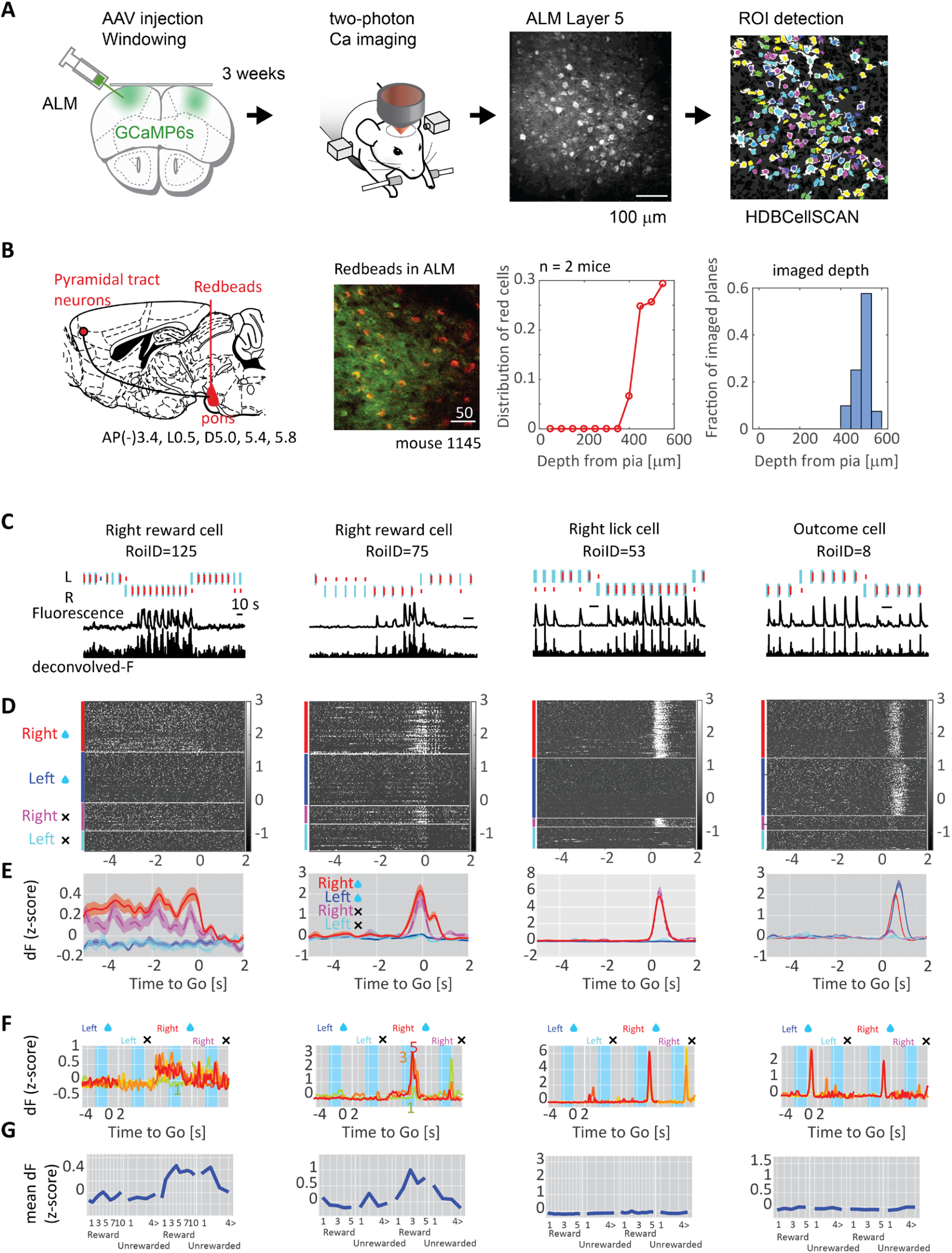
Calcium imaging from ALM layer V cells. (**A**) Procedure of two-photon calcium imaging (Methods). (**B**) Identification of shallow layer Vb neurons in ALM. Pyramidal tract neurons that project to the pons are distributed in the upper layer Vb. Injection of red beads in the pontine nuclei allows us to identify the target depth in our microscope by visualizing the red beads positive cells in ALM. The depth of the imaging plane covered layer Vb of ALM. (**C**) Single-cell examples of fluorescence and deconvolved fluorescence traces during the task. (**D**) Grayscale images showing z-scored activity in four trial types (Right-reward (red), Left-reward (blue), Right-non reward (purple), and Left-non reward (cyan) trials). (**E**) Trial-type averaged activity of (D). Data are mean ± SEM. (**F**) Reward-count or Unrewarded-count averaged activity in the four trial types. The first reward trial means the trial in which the animal will receive the first reward in the state. (**G**) Mean of z-scored activity during the preparatory period (cyan-highlighted region in (F)). X-axis labels showed the cumulative number of rewarded or unrewarded trials within one state.

**Figure S7.**
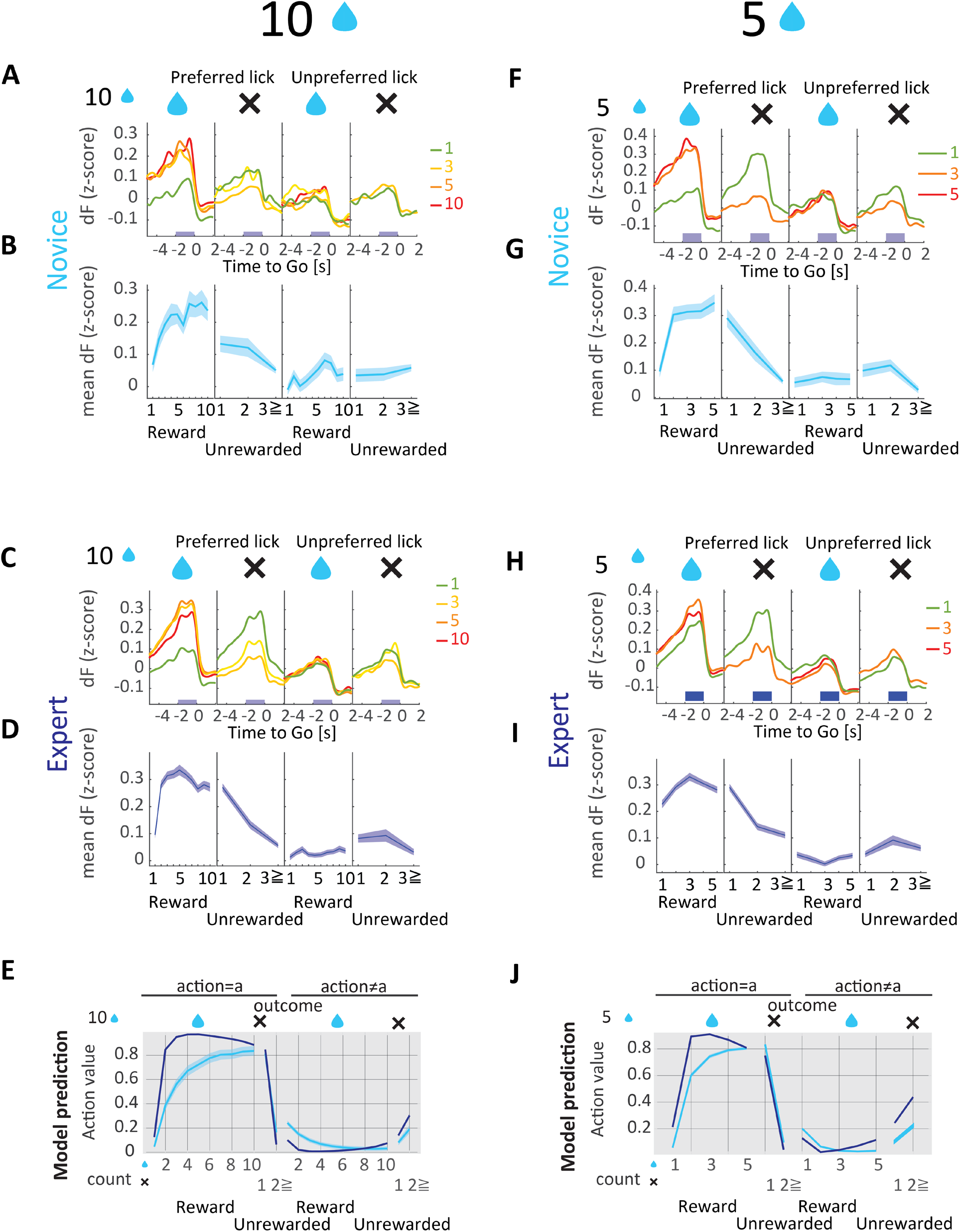
Population activity of preparatory neurons in all four conditions. (**A, C, F, H**) Trial-averaged activity of ramping-up cells that has the peak activity during the preparatory period (Ready to Go, -2 to 0 s to Go cue) in (A) Reward10 novice, (C) Reward10 expert, (F) Reward5 novice, (H) Reward5 expert at various reward counts. Four conditions: Rewarded/Unrewarded × Preferred/Non-Preferred lick trials. The rewarded lick in preferred direction were shown in Fig. 2. (**B, D, G, I**) Average of z-scored activity during the preparatory period by reward and unreward counts within the state. (B) Reward10 novice, (D) Reward10 expert, (G) Reward5 novice, and (I) Reward5 expert. (**E**) Dynamics of action-value predicted by hQ-learning. The meta parameters of hQ-learning were fitted to each session and averaged for plot. In novices, the action-value monotonically increases as a function of reward counts. In experts, the value dynamics develops to a rise-and-decay pattern. Mean ± SEM. (**J**) Same as (E) for Reward5 task.

**Figure S8.**
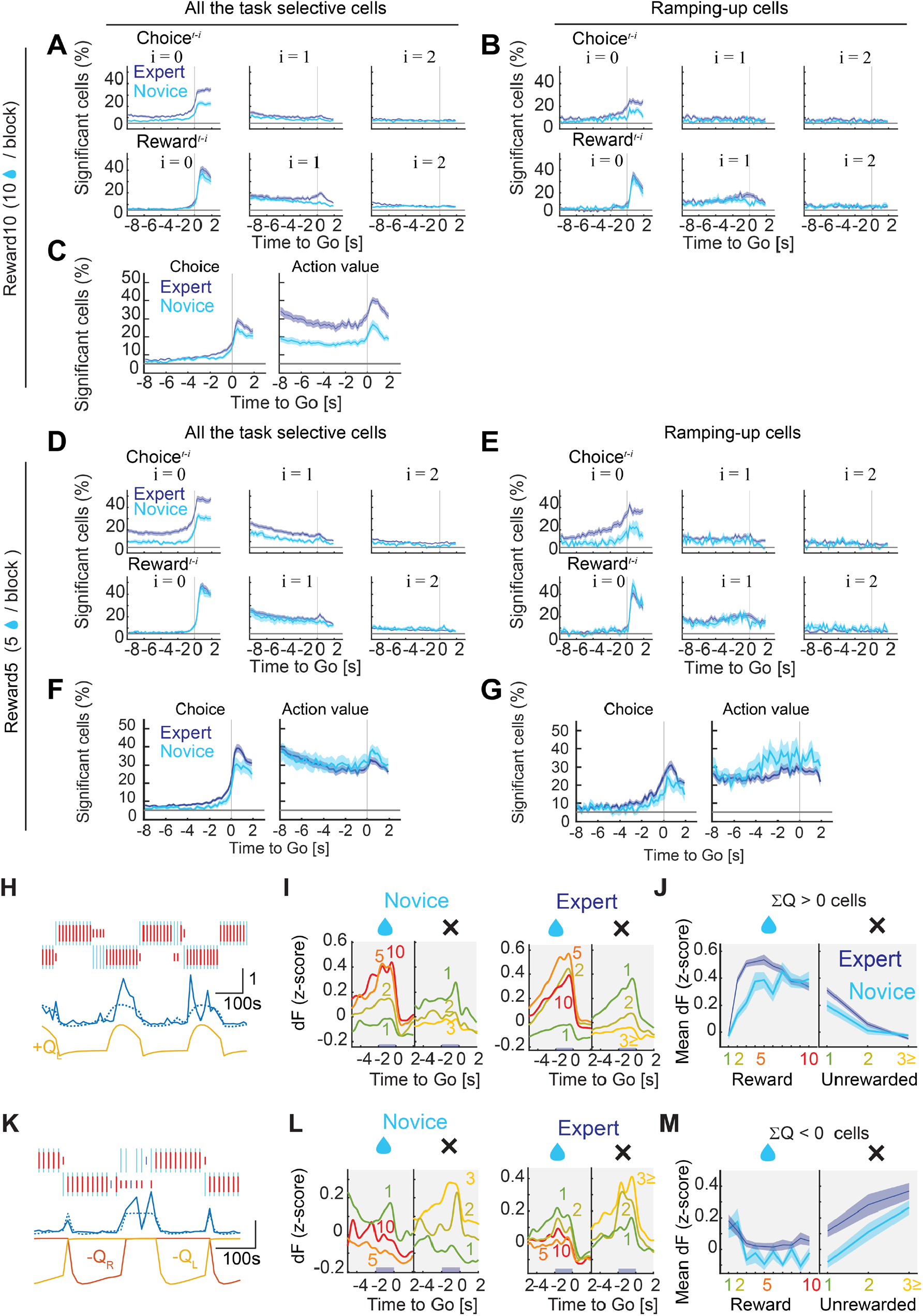
Regression analysis revealed action-value dominance in ALM. (**A**) Percentage of neurons that significantly modulated their activity according to the choice (left or right lick) and reward in the current (i = 0) and previous two (i = 1,2) trials in the non-overlapping 200 ms sliding window (*n* = 4990 cells in expert, *n* = 2889 cells in novice). A task selective cell is defined as the cell with significant p-value (ANOVA) in any of the following time windows; preparatory (−2 to 0 s to the Go cue), action (0 to 1 s), and outcome (1 to 4 s) periods. Reward10 task. (**B**) Same as (A) for ramping-up cells. (**C**) Percentage of cells that significantly changed their activity by choice and action-values computed from all the task selective cells. Choice coding during the preparatory period: novice, 9.4 ± 0.7 %; expert, 11.8 ± 0.6%, *P*_threshold_ = 0.05. Action-value coding during the preparatory period: novice, 17.1 ± 1.5 %; expert, 27.5 ± 1.9 %, *P*_threshold_ = 0.025. (**D**) Same as (A) for Reward5 task. (**E**) Same as (D) for the ramping-up cells in Reward5 task. (**F**) Same as (C) for all task selective cells in Reward5 task. Choice coding during preparatory period; novice, 9.3 ± 1.4 %; expert, 12.8 ± 0.9%, *P*_threshold_ = 0.05. Action-value coding during the preparatory period: novice, 30.2 ± 3.6 %; expert, 27.2 ± 1.8 %, *P*_threshold_ = 0.025. (**G**) Same as (F) for ramping-up cells in Reward5 task. Choice coding during the preparatory period: novice, 10.8 ± 1.8 %; expert, 15.8 ± 1.6%, *P*_threshold_ = 0.05. Action-value coding during the preparatory period: novice, 36.0 ± 6.2 %; expert, 28.3 ± 2.0 %, *P*_threshold_ = 0.025. (**H**) An example of the positive action-value (ΣQ ≥ 0) cell in Reward10 task. The mean z-score of deconvolved fluorescence (dF) during the preparatory period was plotted (blue) overlaid with the predicted value from regression analysis (dashed line). Bottom, the major coefficient (yellow, left action-value Q_L_). (**I**) Trial-averaged activity of positive action-value neurons (defined as cells with significant regressors of Qcontra or Qipsi, and ΣQ ≥ 0) in Reward10 novices and experts. (**J**) Mean of z-scored activity during preparatory period (−2 to 0 s to Go) by reward and unreward counts within the state. Data are pooled from each cell’s preferred directions. (**K**) An example of the negative action-value (ΣQ < 0) cells in Reward10 task. Same format as (H). Bottom, the major coefficients (left and right negative action-values, -Q_L_ and -Q_R_). This neuron showed increased activity following the unrewarded trials. (**L)** Same as (I) for negative action-value cells. (**M**) Same as (J) for negative action-value cells. All mean ± SEM.

**Figure S9.**
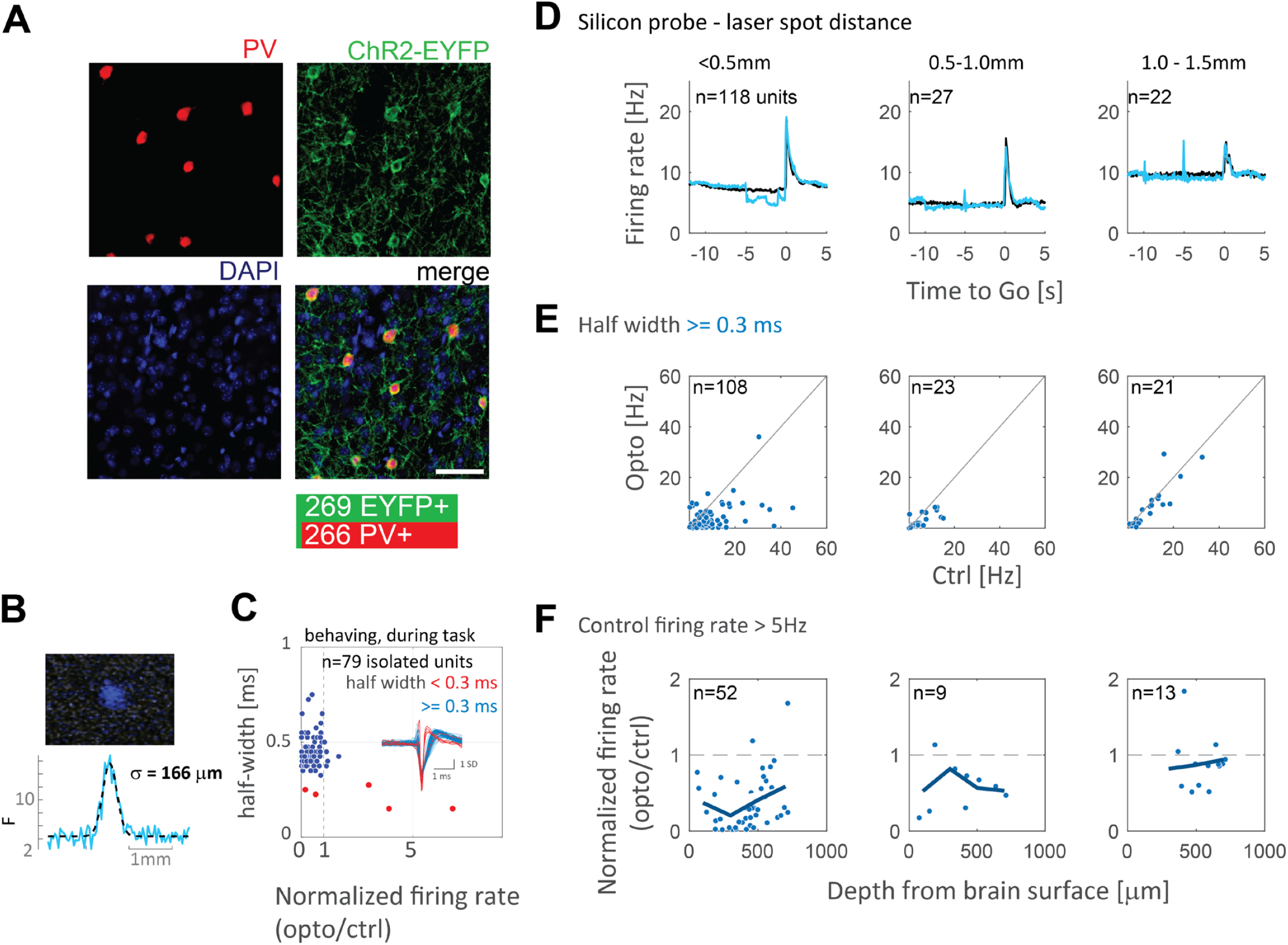
Photoinhibition effects measured in ALM neurons. (**A**) A magnified view showing the expression of ChR2-EYFP in Parvalbumin (PV) positive neurons in ALM of an Ai32 × PV-Cre mouse. Sections were stained with anti-PV and anti-GFP antibodies to visualize PV (red) and ChR2-EYFP (green). Nuclei were counter-stained with DAPI (blue). The majority of ChR2-EYFP positive neurons in the cortex are parvalbumin positive (*n* = 266 out of 269 cells from 2 mice). Scale bar, 50 μm. (**B**) Top, the spatial profile of the laser spot measured at the level of the skull surface. Bottom, the distribution of laser intensity in the lateral distance. Fit with a gaussian distribution indicates that the laser beam has a Gaussian profile with a standard deviation of ∼160 μm. (**C**) Responses of the single-unit activity during the task normalized to the control (stimulation over the head-fixation bar). Each dot corresponds to a single unit. Red circles: units with a half-width less than 0.3 ms (putative inhibitory neurons). Blue circles: units with a half-width broader than or equal to 0.3 ms (putative excitatory neurons). (**D**) Trial-averaged activity in control (black) and photoinhibition (blue) trials. The distance between the laser spot and a silicon probe insertion point was varied from 0 to 1.5 mm. (**E**) Scatter plot of the firing rate in control and photo-inhibited trials. Only the units with half-width ≥ 0.3 msec (putative excitatory neurons) are shown. (**F**) Firing rates normalized to the control trials, plotted as a function of the depth. Only cells with spike rates higher than 5 Hz were analyzed to avoid the floor effect. The depth was calculated from the reading of the micro-manipulator that inserts the silicon probe and the insertion angle.

**Figure S10.**
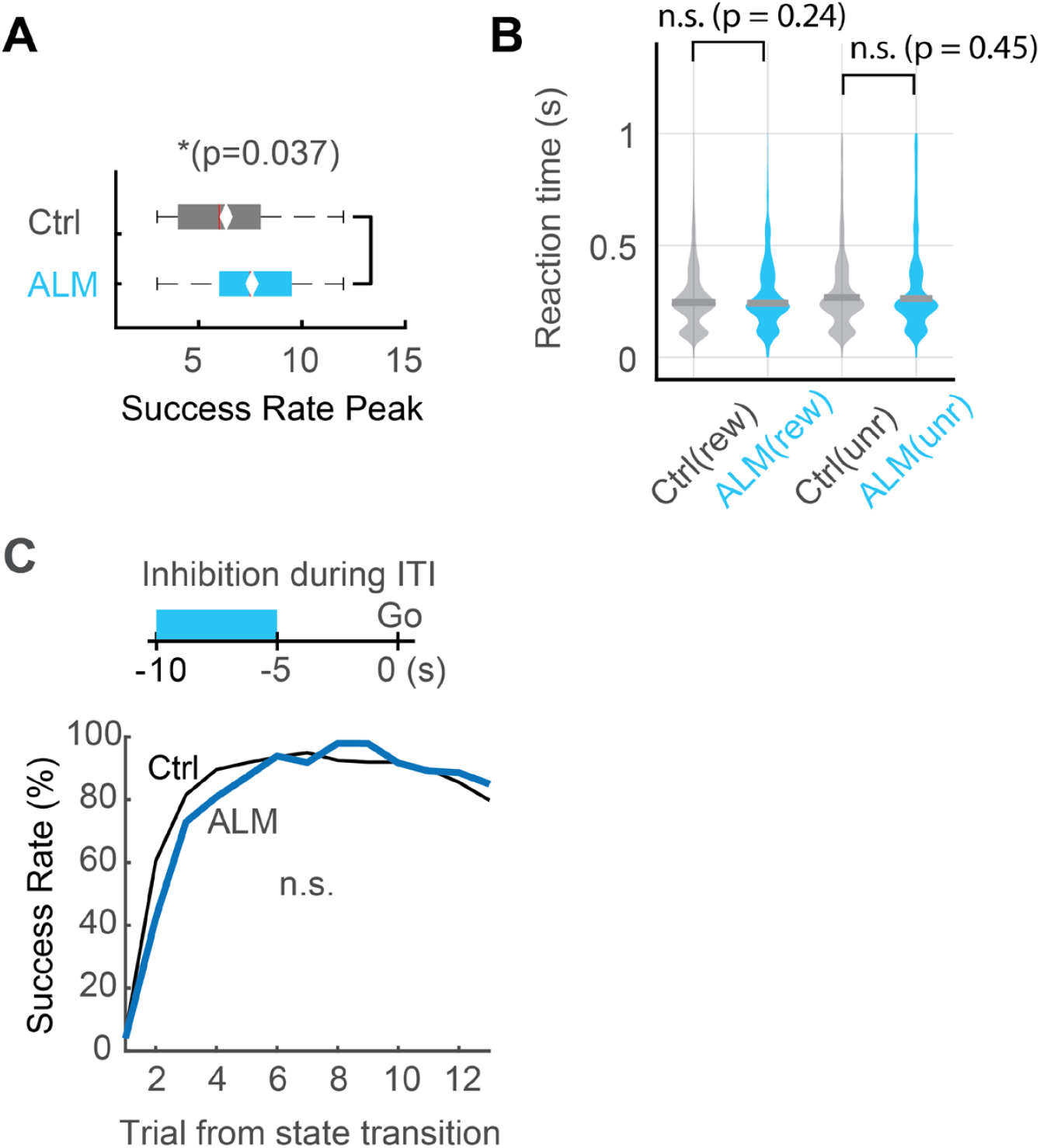
ALM photoinhibition effects on behavior. (**A**) Delayed peak of the success rate in ALM inhibited blocks (Mann-Whitney *U*-test, *\*P* = 0.037). (**B**) Violin plots of reaction time (Go cue to the first lick) before rewarded (rew) or unrewarded (unr) action in control and ALM inhibited trials. No significant difference was detected neither of rewarded nor unrewarded trials (Mann-Whitney *U*-test, rew, *P* = 0.24, unr, 0.45). Ctrl (rew), 245 ± 1.4 ms, *n* = 11850 actions; ALM (rew), 241 ± 3.0 ms, *n* = 3950; Ctrl (unr), 266 ± 12.3 ms, *n* = 3713; ALM (unr), 262 ± 15.6 ms, *n* = 2221. Median ± SEM. (**C**) Photoinhibition during ITI period (−10 to -5 s before the Go cue) had no effects on reversal. Trial-by-trial success rate aligned to the state transition. Control (black) and ALM photo-inhibited (blue) during ITI period (Fisher’s exact test, all *P* > 0.05 with Bonferroni compensation).

**Table S1:** Data preparation. Data filters used to analyze the calcium imaging data set. **1)** Total number of mice, sessions, and cells imaged in each learning stage. **2)**, To eliminate sampling bias, we excluded the same imaging planes within the same learning stage. The same imaging planes can be used for the analysis in different learning stages. **3)**, We then selected cells that showed significantly different responses in any of four trial types (Contra/Ipsi-lick × Rewarded/Unrewarded) during the preparatory period (−2 to 0 s to the Go cue) using ANOVA. **4)**, To analyze the decision-related activity, we selected ramping-up cells that showed maximum activity during the preparatory period in any of the four trial types.

| Learning Stage | Reward10<br>Novice | Reward10<br>Expert | Reward5<br>Novice | Reward5<br>Expert |
| --- | --- | --- | --- | --- |
| **1) Total number of mice, sessions, and cells imaged** | N = 8 mice<br>N = 25 sessions<br>N = 4500 cells | N = 10 mice<br>N = 42 sessions<br>N = 7628 cells | N = 8 mice<br>N = 12 sessions<br>N = 1643 cells | N = 12 mice<br>N = 40 sessions<br>N = 7356 cells |
| **2) After excluding the same image plane** | N = 21 sessions<br>N = 3645 cells | N = 30 sessions<br>N = 5392 cells | N = 11 sessions<br>N = 1561 cells | N = 33 sessions<br>N = 6174 cells |
| **3) Trial-type selective cells (ANOVA)** | N = 965 cells | N = 2741 cells | N = 769 cells | N = 3273 cells |
| **4) Ramping-up cells** | N = 156 cells | N = 459 cells | N = 116 cells | N = 446 cells |

**Supplementary Text1: Reward5 task and short-interval task provides additional supports for reward number dependent behavior in mice**.

To test whether the predictive choice behavior observed in Reward10 task truly depends on the number of rewards, we subjected Rehward10 expert mice to five rewards per choice task (Reward5 task, fig. S2A-C). After additional training in Reward5 task, the expert mice showed the success rate peak before the 5th trial (fig. S2D and E). They also showed a larger increase in win-shift probability (fig. S2F and G) which leads to a higher probability of making reversals at the state transition without mistakes compared to Reward10 experts (movie S2), suggesting that the Reward5 experts learned to make more anticipatory reversals in earlier timing of a state. To exclude the possibility that animals rely on internal timing, the expert mice were also tested in a short inter-trial interval task (from 10 s to 5 s). If a mouse relies on the internal timing to make predictive reversals, reducing the ITI in half would significantly increase the errors, but it did not affect the performance (fig. S3). These results demonstrate that mice can learn the key parameter of the environment, the number of rewards available per state, and predict the state transition triggered by the number of delivered rewards.

**Supplementary Text2: Calcium imaging in Reward5 task further confirmed that ALM value representation depends on the number of available rewards in a state**.

To investigate the development of value representation in a different task parameter (here, Reward5), we imaged ALM layer V cells of Reward5 novice and expert mice (*n* = 6047 cells from 44 sessions, 14 mice are analyzed. *n* = 116 in novice and 446 cells in expert were classified as ramping-up cells). As we observed in Reward10 task, a significant number of ALM layer V cells modulated their activity by the future choice and reward history (for single cell example, see fig. S6: for regression analysis, see fig. 8D and E). Indeed, the dynamics of population preparatory activity reverted to a monotonic increase in novices and then evolved into the rise-and-decay dynamics again in experts (fig. S7F-I) which is consistent with the model prediction (fig. S7J). Most importantly, the value dynamics was compressed in Reward5 experts compared to Reward10 experts (fig. S7G and I), suggesting that the dynamics of preparatory activity depends on the number of available rewards in one state, and not the innate tendency to avoid repeating the same action. The regression analysis using the choice and action values further confirmed that ALM layer V is enriched with the action value coding cells (fig. S8 F and G). These imaging results further confirmed our key finding that ALM preparatory neurons encode the prospective value computed from the knowledge of the environment.

**Supplemental Movie1: ALM layer V calcium imaging**.

Imaged at Anterior 2.5 mm from bregma, Left 1.5 mm, 500 μm deep. x8 speed, GCaMP6s.

**Supplemental Movie 2**: **Reward5 expert behavior**.

An example of Reward5 expert behavior in which the animal is making reversals without mistakes.

